# An Improved Algorithm for Inferring Mutational Parameters from Bar-Seq Evolution Experiments

**DOI:** 10.1101/2022.09.25.509409

**Authors:** Fangfei Li, Aditya Mahadevan, Gavin Sherlock

## Abstract

Genetic barcoding provides a high-throughput way to simultaneously track the frequencies of large numbers of competing and evolving microbial lineages. However making inferences about the character of the evolution that is taking place remains a difficult task. Here we describe a step toward more accurate inference of fitness effects and establishment times of beneficial mutations, which builds upon a prior method by enforcing self-consistency between the population mean fitness and the individual effects of mutations within lineages. By testing our inference method on a simulation of 40,000 barcoded lineages evolving in serial batch culture, we find that this new method outperforms its predecessor, identifying more adaptive mutations and more accurately inferring their mutational parameters. We have made available our code for the joint Bayesian inference of population mean fitness and lineage-specific mutational parameters, in the hope that it can find broader use by the microbial evolution community.

## Introduction

Clonal interference is a phenomenon that occurs in large asexual populations, in which multiple beneficial mutations arise contemporaneously and compete with each other without recombining onto the same genetic background (Gerrish and Lenski, 1998; Desai and Fisher, 2007; Park and Krug, 2007; Fogle et al., 2008; Lang et al., 2011). Although these mutations may later accumulate onto the same genomes due to high mutation rate (Bollback and Huelsenbeck, 2007), clonal interference remains an important evolutionary force across a wide range of timescales. Experimental evolution in large microbial populations, where the emergence of beneficial mutations is common enough that clonal interference is widespread, has been widely used to explore this regime of adaptation (Lenski et al., 1991; Miralles et al., 1999; de Visser and Rozen, 2006; Kao and Sherlock, 2008; Good et al., 2017). In such experiments, microbes such as fungi, bacteria, or viruses are propagated for hundreds (or thousands) of generations in a controlled experimental system, typically either by serial transfer of batch cultures or continuous culture. Adaptive mutations that emerge during such an experiment can be identified by whole-genome sequencing (WGS) of multiple isolates from the evolved populations (Araya et al., 2010; Kvitek and Sherlock, 2011; Tenaillon et al., 2012; Hong and Gresham, 2014).

However, this approach has limitations. Firstly, it cannot provide information about the occurrence time and fitness effect of mutations. Secondly, it can only identify the subset of adaptive mutations that reaches high frequency, which tends to consist of those which arose earlier or provide a larger fitness benefit. Although the minimum frequency at which the mutation is detectable can be lowered by sequencing more isolates, the high cost of WGS (in comparison to amplicon sequencing) quickly makes this method impractical for identifying low-frequency mutations. An alternative to WGS for multiple isolates at the end of an evolution experiment is to conduct WGS for the whole population at multiple time points during the evolution (Kvitek and Sherlock, 2013; Lang et al., 2013; Good et al., 2017). With such a time series, one can roughly estimate the occurrence time of some of the mutations. However, this method is more expensive, does not provide the fitness effects of mutations, and fails to identify mutations at a frequency lower than ~ 1% under typical financial constraints.

To better explore these low-frequency dynamics, a high-resolution lineage tracking system was developed in *S. cerevisiae*, based on a genetic barcoding platform (Levy et al., 2015). This system is capable of detecting thousands of initial adaptive mutations in an originally clonal evolving population, by simultaneously monitoring the relative frequencies of ~ 5 × 10^5^ lineages, each defined by a unique genetic barcode of 20 nucleotides and consisting of ~ 100 cells initially in a typical experiment (Levy et al., 2015). With the high-resolution information on lineage frequencies from the barcode counts at multiple time points, one can estimate fitness effects and establishment times of adaptive mutations using a statistical framework based on the theory of branching processes and Bayesian inference (Levy et al., 2015). Here we refer to this algorithm as `FitMut1`. `FitMut1` can detect adaptive mutations at frequencies higher than ~ 10^-6^ from barcode frequencies over time, and can be followed by WGS for clones with different barcodes for further characterization of mutations at the genotypic level. For this step, isolating clones is relatively straightforward since each lineage contains an unique barcode that can be easily recognized by Sanger sequencing (Venkataram et al., 2016). In addition to *S. cerevisiae*, other microbes such as *E. coli* have been studied with similar barcoding approaches (Jahn et al., 2018; Jasinska et al., 2020).

It should be emphasized that not all beneficial mutations are detectable. A minimum requirement for a mutation to be detected is its *establishment* (Desai and Fisher, 2007; Levy et al., 2015). For a beneficial mutation that occurs initially in a single cell and with fitness effect *s*, there is a substantial probability of going extinct soon after occurring, due to random fluctuations, even though the mutant confers a growth advantage. However, if a mutant gets “lucky enough” (with the probability proportional to *s*) to reach a certain size (proportional to 1/*s*), it will grow exponentially with rate *s* thereafter. In this case, we say that the mutation carried by the mutant has *established*. By extrapolating its exponential growth backward in time until the mutant population crosses the rough boundary between stochastic and deterministic dynamics, we can define an *establishment time* as the time after which the mutant cells effectively grows deterministically. Establishment time roughly reflects the occurrence time of a mutation, up to uncertainty on the order of 1*/s*.

Nevertheless, the exponential growth rate of an adaptive lineage (in which an adaptive mutation has established) cannot be measured directly to yield the fitness effect of the mutation. This is because 1) the mutation must sweep through the entire lineage before dynamics of the lineage reflect those of the mutation, and 2) the lineage trajectory in a well-mixed environment bends over as it competes against the increasing population mean fitness. Therefore, the mean fitness is required for accurately characterizing the dynamics of adaptive lineages and further inferring establishment times and fitness effects of mutations. However, it is difficult to measure the mean fitness directly. In `FitMut1`, the mean fitness is estimated by monitoring the decrease in frequency of neutral lineages (those without an established mutation) between consecutive sequencing time points. However, `FitMut1` fails when the number of available neutral lineages is insufficient, which can happen when the sequencing read depth is low, the bottleneck size (number of cells per barcode at the bottleneck) is small, or the mean fitness increases rapidly.

In this work, we describe an improved algorithm, `FitMut2`, which iteratively estimates the mean fitness and characterizes the adaptive lineages in an evolving population, without relying on the number of available neutral lineages. This makes `FitMut2` less impaired by low sequencing read depth, small bottleneck size, or rapidly increasing mean fitness, and thus more robust and accurate than `FitMut1`. To assess the performances of both algorithms, we ran `FitMut1` and `FitMut2` on the same simulated dataset and compared their outputs with the ground truth.

We first introduce `FitMut2`, which includes a summary of `FitMut1` and the modifications that constitute `FitMut2` (Methods:A). In addition, we also discuss the simulated data that we used to benchmark the performance of `FitMut2` and compare it to `FitMut1` (Methods:B). We then evaluate the performance of `FitMut2` on simulated data and compare it with `FitMut1` on the same dataset (Results). Finally, we discuss the limitations of `FitMut2` and possible future improvements (Discussion).

## Methods

### A. Algorithm Overview

`FitMut1` models the dynamics of lineage growth with a stochastic branching process, thereby associating with each lineage a probability distribution of the number of reads that map to its barcode, conditional on this read number at a previous time point. In addition to the demographic stochasticity of births and deaths, this distribution considers various sources of noise: cell transfer, DNA extraction, PCR and sequencing, which are represented by a phenomenological parameter *κ_k_* for each time point *t_k_* (details in S4). Our noise model is consistent with a branching process, wherein (conditional on the read number at *t*_*k*–1_) the variance in read number at *t_k_* is proportional to the mean read number at *t_k_*, with constant of proportionality 2*κ_k_*. In `FitMut1`, first the mean fitness of the population 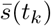 and the noise parameter *κ_k_* are estimated for all sequencing time points *t_k_* by monitoring the decline of neutral lineages, which are assumed to constitute the majority of lineages at intermediate read number. Assuming that each lineage contains a mutation with fitness effect *s* and establishment time *τ*, the posterior likelihood of each lineage trajectory data is calculated, and maximized over *s* and *τ* to find the values for which the observed data are most likely. `FitMut1` relies on neutral lineages for estimating the mean fitness, which then influences all subsequent steps. Although this works well when the number of neutral lineages is large (e.g. at the beginning of a typical experiment), the number of neutral lineages falls dramatically during the evolution as beneficial mutations increase in frequency in the population, and the efficacy of `FitMut1` thus decreases.

Instead of relying on neutral lineages, `FitMut2` uses an iterative approach to self-consistently infer the population mean fitness together with *s* and *τ* for each putative mutation. This approach does not require a large number of neutral lineages to be present, and enforces that the individual mutations and their frequencies are consistent with the inferred population mean fitness. `FitMut2` only relies on the number of putatively neutral lineages to estimate the noise parameter *κ_k_* at each sequencing time point, and we have found that the inference results are not very sensitive to the value of this parameter. With this self consistent method, `FitMut2` identifies more adaptive mutations and obtains the *probability* of a lineage being adaptive conditional on the data. By contrast, `FitMut1` provides a ratio of posterior likelihoods, which is not required to be between 0 and 1, and is harder to interpret. The algorithm of `FitMut2` proceeds as follows:

1. For each sequenced time point: Initialize the mean fitness to 0 and calculate *κ_k_* from the empirical distribution of read numbers assigned to putatively neutral lineages at that time.
2. For each lineage: Use Bayes’ theorem to calculate the probability that the lineage is adaptive given the observed read number trajectory, under a prior distribution over fitness effect *s* and establishment time *τ* (the choice of prior is discussed further in the Discussion). If the probability is greater than 0.5, designate the lineage adaptive, and maximize the posterior likelihood over *s* and *τ* to find the most likely fitness effect and establishment time under the assumed prior (see details in S5).
3. Estimate the number of mutant cells over time for each identified adaptive lineage, using the inferred fitness effect and establishment time of the mutation. Update the mean fitness accordingly.
4. Repeat steps 2 and 3 until the estimated mean fitness at each sequencing time point converges to a self-consistent value.

By design, the mean fitness 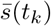 at time point *t_k_* must agree with the mean fitness from the adaptive lineages and their frequencies, given by 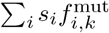 where *s_i_* is the fitness effect of the mutation of lineage *i* and 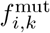 is the frequency of mutant cells at time point *t_k_*. Although these two ways of estimating the mean fitness are shown to roughly agree in `FitMut1` (Levy et al., 2015), their equality is explicitly enforced here, and it improves the algorithm’s accuracy when the read depth is low or the mean fitness is rapidly increasing.

### B. Simulation

Numerical simulation is an effective method to evaluate performance of the algorithm when available experimental data are limited. Here, we evaluated the performance of `FitMut2` using a simulated dataset, which allows us to compare the inference result with the ground truth. Our numerical simulations consider the entire process of a barcode-sequencing (bar-seq) evolution experiment of a barcoded cell population using serial batch cultures (Figure 1). Starting from a single cell per barcode, lineages first go through 16 generations of pregrowth without competition, during which mutations can and do occur. This step simulates the cell growth on agar plates after the barcode transformation but before the evolution experiment begins. Each cell into which a barcode was successfully transformed can grow into a colony on an agar plate, with all cells in the colony containing the same barcode. The pregrowth phase includes two processes which are inevitable in the experimental process of building a barcode library: 1) the growth noise on agar plates, which generates a non-uniform distribution of lineage sizes and 2) the occurrence of mutations before the evolution experiment commences. Both of these features can significantly influence the evolutionary dynamics. After being scraped from agar plates, cells of the colonies are pooled together and grown up overnight before being sampled and inoculated into the medium. In the simulation, we ignore this process, because it includes very few generations of growth.

**Figure 1.**
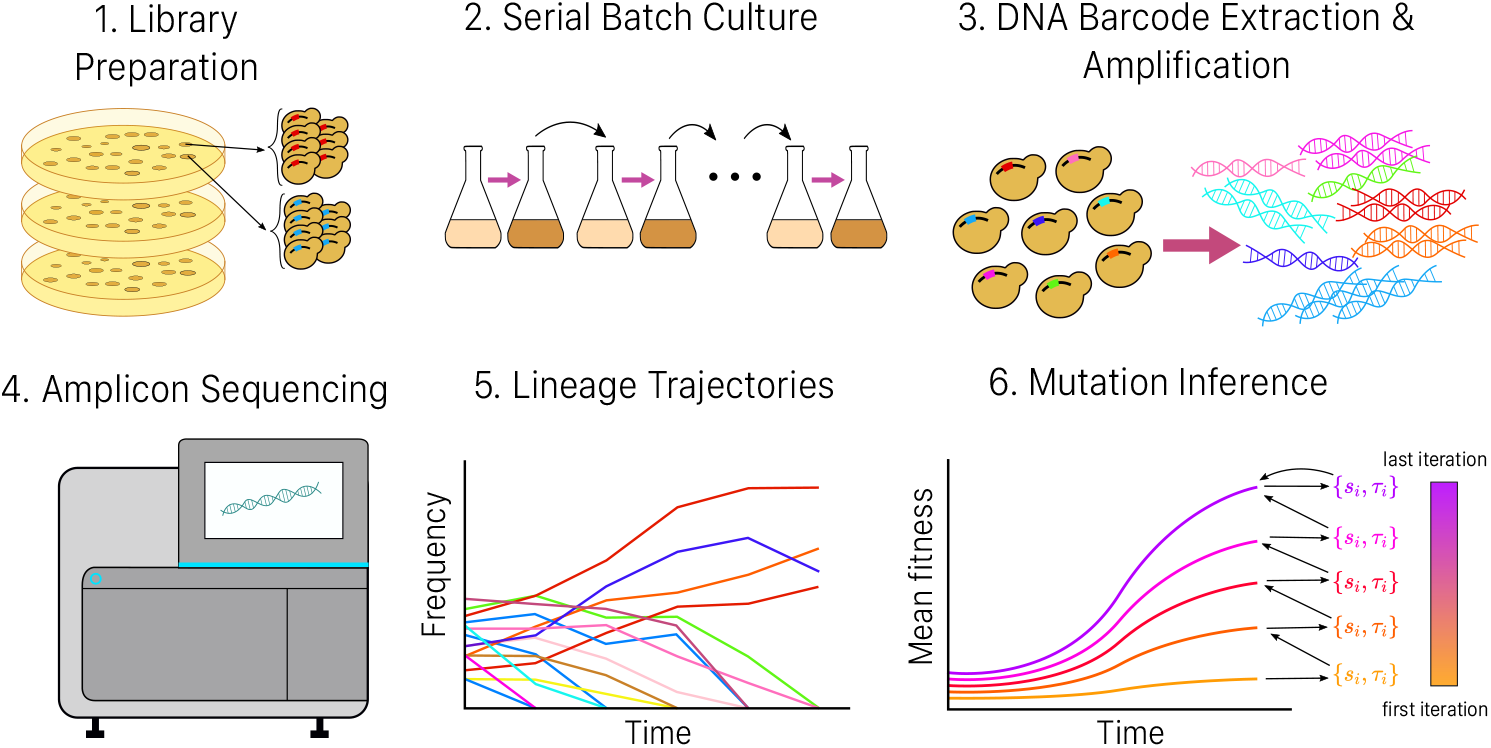
Procedure of a complete barcoded evolution experiment with analysis included. Steps 1 to 4 depict the procedure of a typical bar-seq evolution experiment, which gives rise to a series of lineage trajectories over the course of the experiment (Step 5), with each lineage defined by one barcode. Different colors represent lineages with different barcodes. Step 6 is a schematic of how we use these trajectories to identify adaptive mutations that occurred in the evolution experiment, and self-consistently infer the fitness effects and establishment times of individual mutations together with the mean fitness of the population (Methods:A).

To initialize the evolution experiment with a barcode library consisting of *L* unique barcodes, 100*L* cells are sampled from the population after the pregrowth. This yields a mean of 100 cells per barcode, which roughly corresponds to parameters commonly used in an experiment. The barcoded population is then evolved through pooled growth by serial batch culture. Each growth cycle consists of *g* generations of stochastic doubling, after which a fraction 1/2^*g*^ of the cells from the end of the growth cycle is sampled and transferred to another fresh culture. This process of growth and dilution, with *B* cells transferred at each bottleneck, can be thought of as constant-population process with an *effective population size* given by *gB* and a per-generation offspring number variance 2*c* ≈ 2, as discussed in Section S3. This effective description is very useful for quantitatively matching theory to experiment, and is essential for the functioning of both `FitMut2` and `FitMut1`. Hereafter, we use the term *effective lineage size* to refer to the lineage or population size that would be necessary to obtain the same statistics of lineage fluctuations if the total number of cells were constant in time rather than growing by a factor of 2^*g*^ every cycle.

Although our simulations can keep track of an arbitrary number of mutations per cell, we have not pursued the inference of these later mutational effects in the current work. Instead we make the simplifying hypothesis that at most one beneficial mutation occurs per cell. In light of evidence suggesting that the distribution of fitness effects (DFE) of the second mutation in a cell is different from that of the first mutation, due to epistasic or physiological constraints (Aggeli et al., 2021), this hypothesis allows us to focus on *initial* adaptive mutations. Although each simulated individual can obtain at most one beneficial mutation, only ~ 3% of lineages contain more than one established mutation in our simulations; when this occurs we record the “true” fitness of the lineage as the *s* of the mutation that generates the maximum number of mutant cells by the end of the evolution. A mutation that occurred with fitness effect *s* is counted as established if, at any time during the evolution, the mutant’s instantaneous frequency reaches 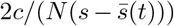. We have found that typically, on the order of 20 of the highest fitness mutations are sufficient to account for the majority of the mean fitness increase over the simulation. However we are able to identify many more mutations than these, though they do not contribute substantially to the mean fitness.

At each cell-transfer time point over the course of the experiment, 500*L* cells are sampled from the saturated population to simulate the process of genome DNA extraction, and go through 25 rounds of stochastic doubling, to simulate PCR with 25 cycles (a larger number of cycles will not have a significantly larger effect on the PCR noise, since only the initial doublings contribute to the stochasticity). Then an extra sampling of the size *rL* after PCR is performed to simulate the noise introduced by sequencing, with *r* being the average sequencing read number per lineage per time point.

The entire process generates a lineage trajectory over time for each barcode. The simulation includes five potential sources of noise: cell growth, sampling during cell transfers, DNA extraction, PCR, and sequencing. Each step is modeled by a layer of Poisson noise (including for each generation of cell growth and each cycle of PCR). To test the performance of `FitMut2`, we ran simulations with four different underlying DFEs denoted by *μ*(*s*), where *μ*(*s*)*ds* is the rate of mutations with fitness effect in the interval (*s,s* + *ds*) (details of the DFEs we simulated are in Section S9). The total beneficial mutation rate is given by 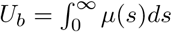. For each simulation, a population of 4 × 10^4^ single cells undergoes 16 generations of pregrowth before all these lineages are pooled and grown by serial batch culture for *T* = 112 generations, with *g* = 8. For each of four DFEs, sequencing is simulated with four different average read numbers per lineage *r* = 10,20,50,100.

Inference is performed with the prior distribution for all conditions: 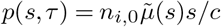. Note that ∫*p*(*s,τ*)*dsdτ* is approximately the number of established mutations per lineage. The factor of *s/c* arises from establishment probability ~ *s/c* in the branching process model, and *n*_*i*0_ is the effective size of lineage *i* at *t*_0_. 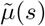 is the prior we take for the DFE, which is 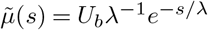 with *λ* = 0.1 and *U_b_* = 10^-5^ throughout this paper.

Figure S1 shows trajectories of all lineages in one of our simulations with an exponential DFE and *r* = 100, corresponding to the 1st row and 4th column in Figure 2.

**Figure 2.**
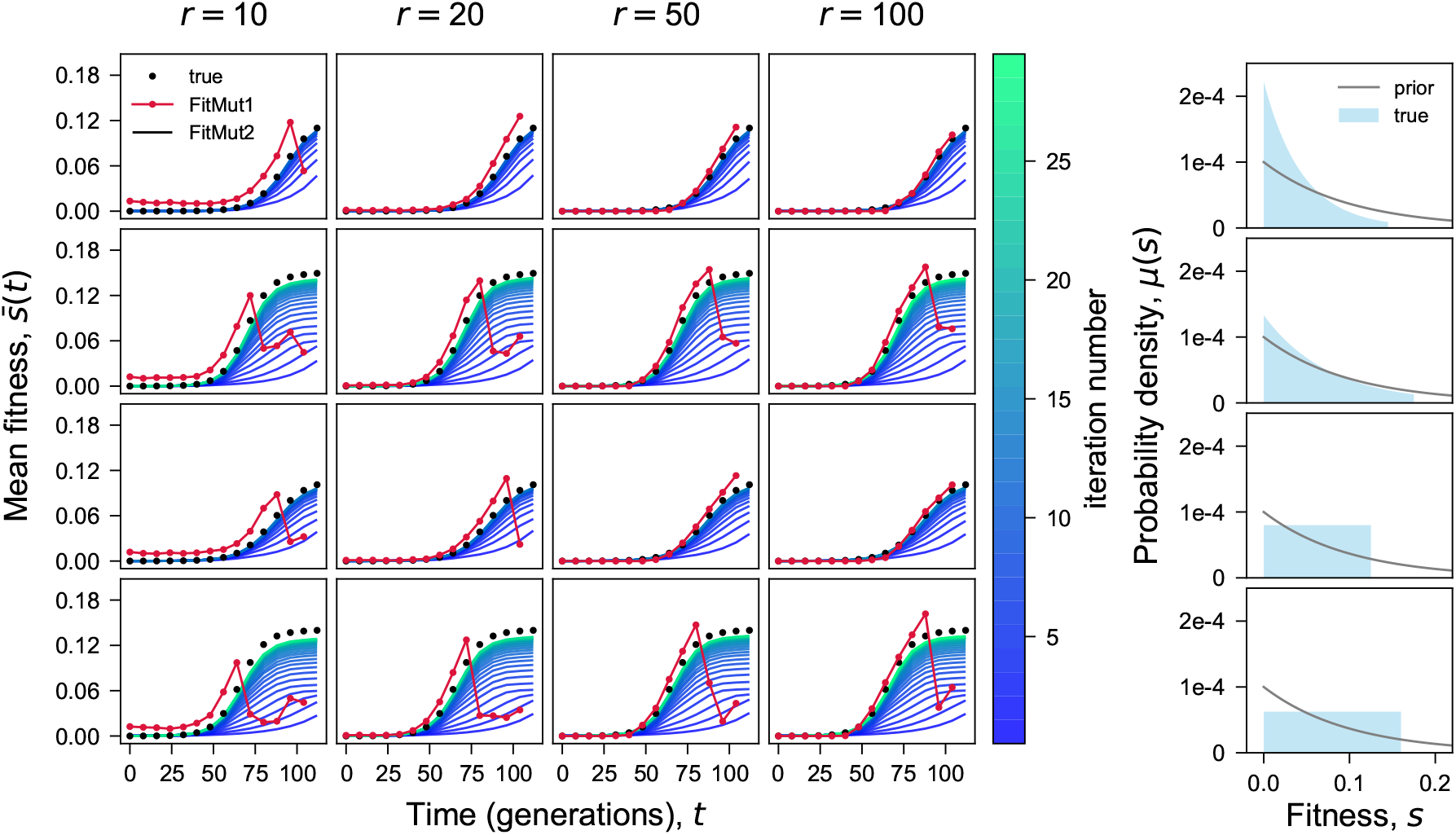
Iterative inference of the mean fitness. Comparison of the true mean fitness 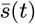 with the mean fitness inferred by both `FitMut1` and `FitMut2`, for different sequencing depths (columns) and *μ*(*s*) (rows). Each row in the 4 × 4 array corresponds to one simulation of the evolution, with the columns differing by the average simulated sequencing depth per time point per lineage. The 5th column shows the DFE *μ*(*s*) used in each simulation condition, and the prior 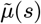 used for both `FitMut2` and `FitMut1` across all simulated DFEs.

## Results

### A. Fitness per generation vs. fitness per cycle

Before presenting the results of running `FitMut2` on our simulated dataset, we discuss an important aspect of interpretation that should be kept in mind whenever one is analyzing data from serial batch culture. How should we interpret the parameters output by our inference algorithm? Specifically, what is the meaning of the fitness increment *s* obtained by a particular lineage in a biological sense? Previous work has explored the importance of variable growth conditions *over a single cycle of batch culture* in creating a much more complex environment than meets the eye (Li et al., 2018). In particular, selection pressure varies over a single cycle of batch culture as the environment, created by the time-varying yeast (or any other organism) metabolism, shifts dramatically. Within a single cycle of batch culture spanning *g* generations, the relevant quantity for the evolutionary dynamics is not *fitness per generation*, but rather *fitness per cycle*. In fact, many adaptive mutations may have time-varying fitness effects over a single cycle. In *S. cerevisiae*, which undergoes multiple metabolic shifts over a single cycle, a beneficial mutation can be neutral during the first seven generations (fermentation), but have a large fitness advantage during the last generation of the cycle (respiration). Therefore it is misleading to interpret results in terms of a fitness per generation: we cannot define a singular notion of “fitness” without accounting for the interaction between organism and environment. One should avoid claiming more granularity than one’s highest temporal resolution, which here is the length of a single batch culture cycle. Since the theory on which `FitMut2` is based considers a simpler effective model without time-varying fitness, fitness effect per generation is used in the branching process model we use. However, the trouble arises if the theory is taken too seriously in interpreting experimental results. Although we report results in terms of fitness per generation in our inference algorithm for both the population mean 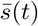 and for adaptive mutation *s*, we emphasize that in real experimental conditions there is little reason to believe that these values correspond to anything other than an *average* quantity that depends on experimental setup and conditions.

### B. FitMut2 robustly estimates mean fitness

`FitMut2` and `FitMut1` differ essentially in how they infer the population mean fitness — this then leads to differences in the inference of mutational parameters. Figure 2 shows the mean fitness trajectories inferred by both `FitMut2` and `FitMut1`. The number of iterations required to converge upon a self-consistent mean fitness is larger for simulations with a wider DFE, but rarely exceeds 40, and convergence appears monotonic. `FitMut1` estimates the mean fitness accurately when sequencing read depth is high (i.e. *r* = 100), and the mean fitness increases slowly (all time points for DFEs with small variance, or early time points for DFEs with large variance). However, for low sequencing read depth (*r* = 10 or 20), or as the mean fitness increases rapidly (later time points for DFEs with large variance), `FitMut1` begins to perform poorly.

### C. FitMut2 accurately estimates mutational parameters

We examined how accurately `FitMut2` estimates fitness effects and establishment times by comparing its inferences to the truth from our simulated dataset (Figure 3). While numerous adaptive mutations are not detected by either algorithm, `FitMut2` identifies hundreds of adaptive mutations missed by `FitMut1` at low read number *r* = 10 and a wide DFE (Results:D), while maintaining a negligible false positive rate (Figure 5B). For adaptive mutations detected by each algorithm, we compare inferred values of parameters to the truth in the simulation. For the adaptive mutations detected by `FitMut2`, there is a very strong correlation between the true fitness effect and the inferred value, and fairly strong correlation between the true occurrence times and the inferred establishment times. For comparison, we also show the results from `FitMut1` in Figure S2, and we see that our new algorithm significantly outperforms the old algorithm. To further assess inference accuracy for mutations identified as adaptive by both `FitMut2` and `FitMut1`, we compared the estimation error between `FitMut2` and `FitMut1` (Figure 4). `FitMut2` has improved accuracy over `FitMut1`, particularly for those simulations in which `FitMut1` could not estimate the mean fitness accurately. For the simulations in which `FitMut1` underestimates mean fitness, the fitness effects and establishment times are also underestimated, which affects the subsequent estimation of the DFE (see the 5th column in Figures 3A and S2A).

**Figure 3.**
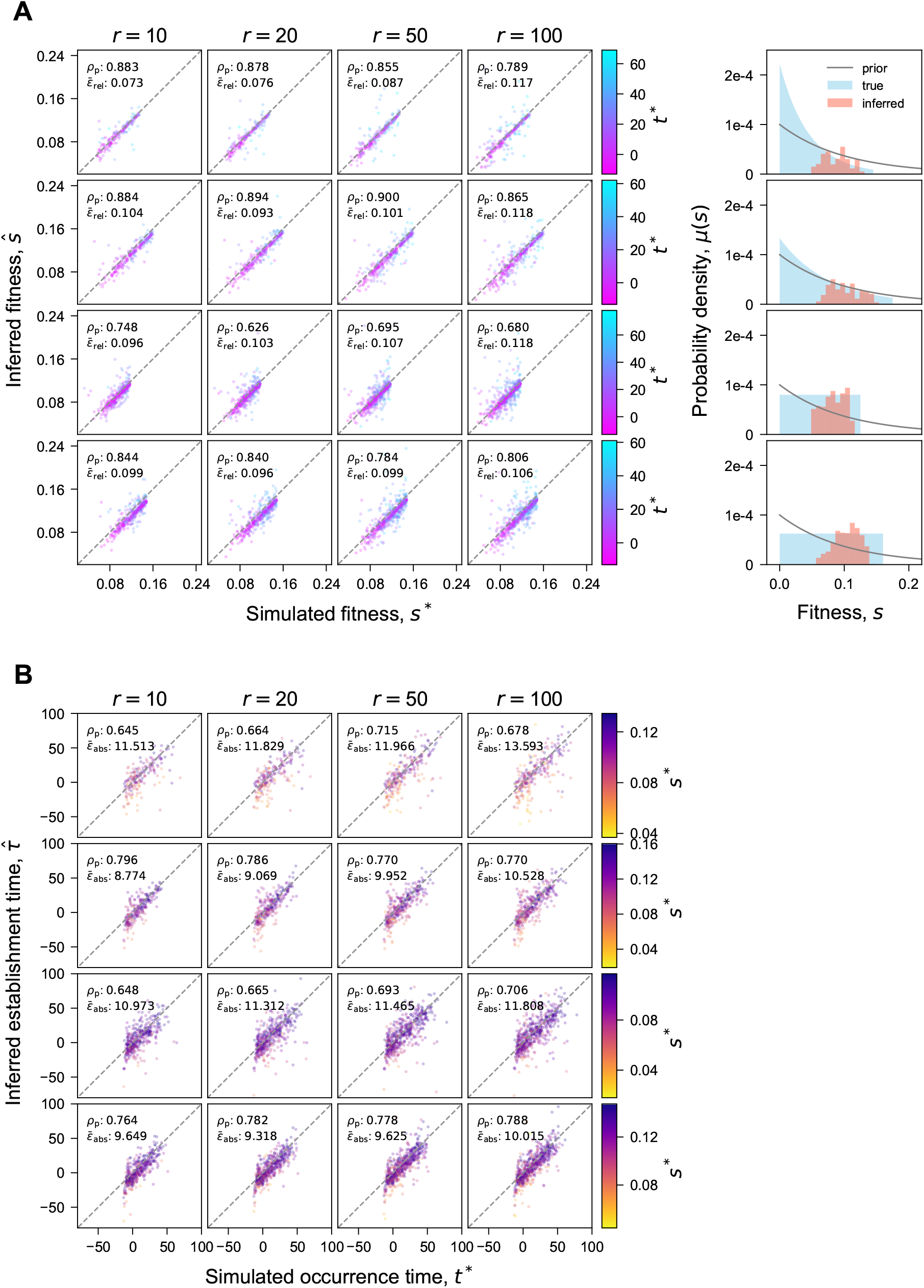
Inference accuracy of (A) fitness effects and (B) establishment times (FitMut2). Comparison of inferred *s* and *τ* with simulation. Each panel in the 4 × 4 array corresponds to one simulation (Methods:B). Each point is an adaptive mutation that established in the simulation and was identified by `FitMut2`. Points in (A) are colored by their true occurrence time *t*\*, while points in (B) are colored by their true fitness *s*\*. Negative *t*\* indicates adaptive mutations that occurred during pregrowth. *ϵ*_rel_ in (A) is defined as 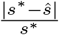 and *ϵ*_abs_ in (B) is defined as 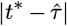. *ρ_p_* is the Pearson correlation coefficient. The 5th column in (A) shows the comparison between *μ*(*s*) and the inferred DFE (estimated as in S8). In Figure S2 we show the same data for `FitMut1`.

**Figure 4.**
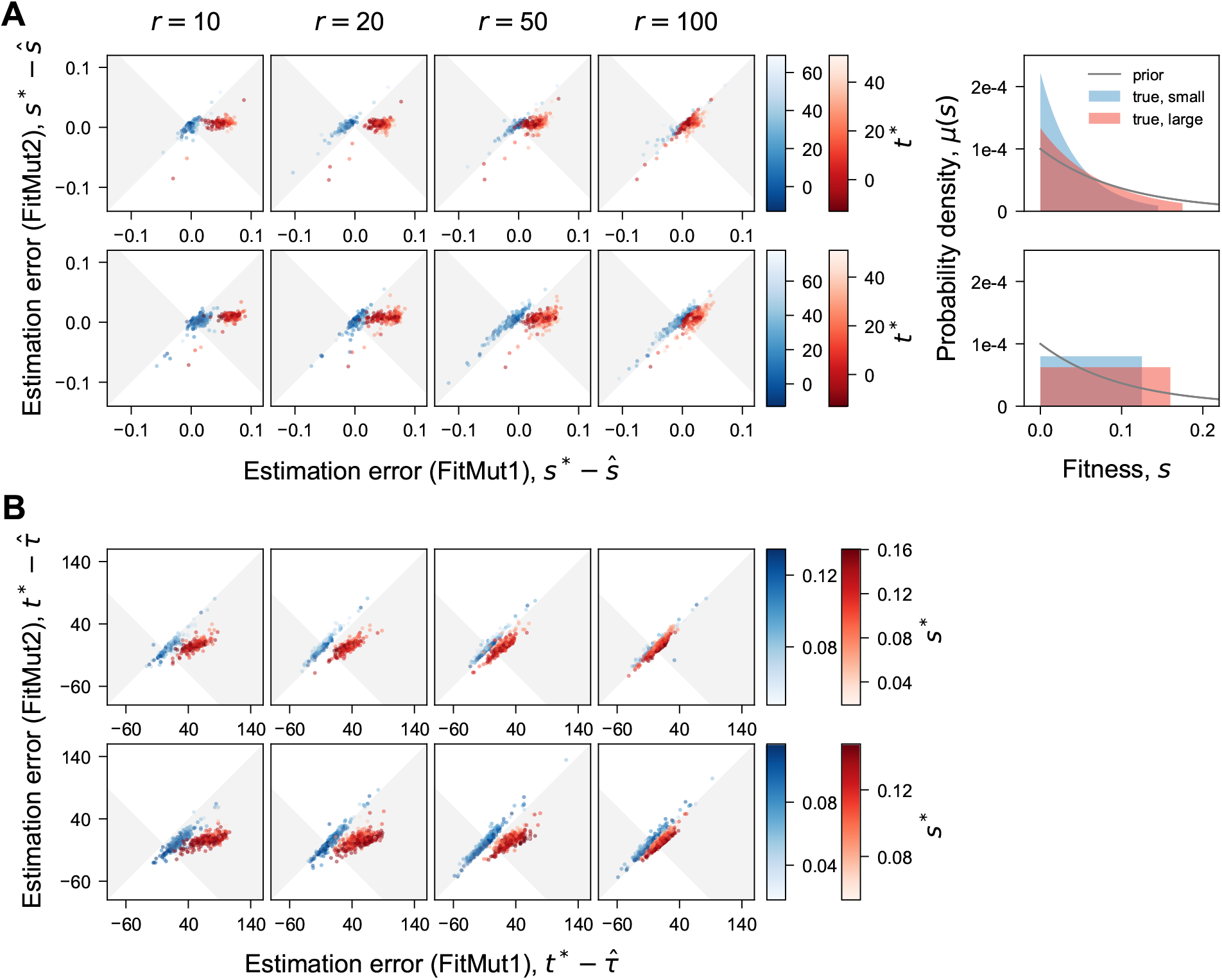
Estimation error of the fitness effects and the establishment times. Comparison of the estimation error, measured between simulation and inference, for `FitMut1` and `FitMut2`. (A) shows fitness effects and (B) shows establishment times. Each column corresponds to an average number of reads per lineage *r*. Different rows correspond to different classes DFEs: exponential and uniform. Each panel includes the inference error of two simulations (Methods:B) from the same family of *μ*(*s*) with different variances (blue for smaller variance, red for larger variance). Each dot in the scatter plot represents an adaptive mutation that established in the simulation and was identified by both `FitMut2` and `FitMut1`. Dots falling within the gray region indicate the adaptive mutations that were more accurately inferred with `FitMut2` than with `FitMut1`. Blue and red dots are colored by their occurrence time *t*\* for (A), and by their true fitness *s*\* for (B). *μ*(*s*) is plotted in the 5th column (blue for small variance, red for large variance).

**Figure 5.**
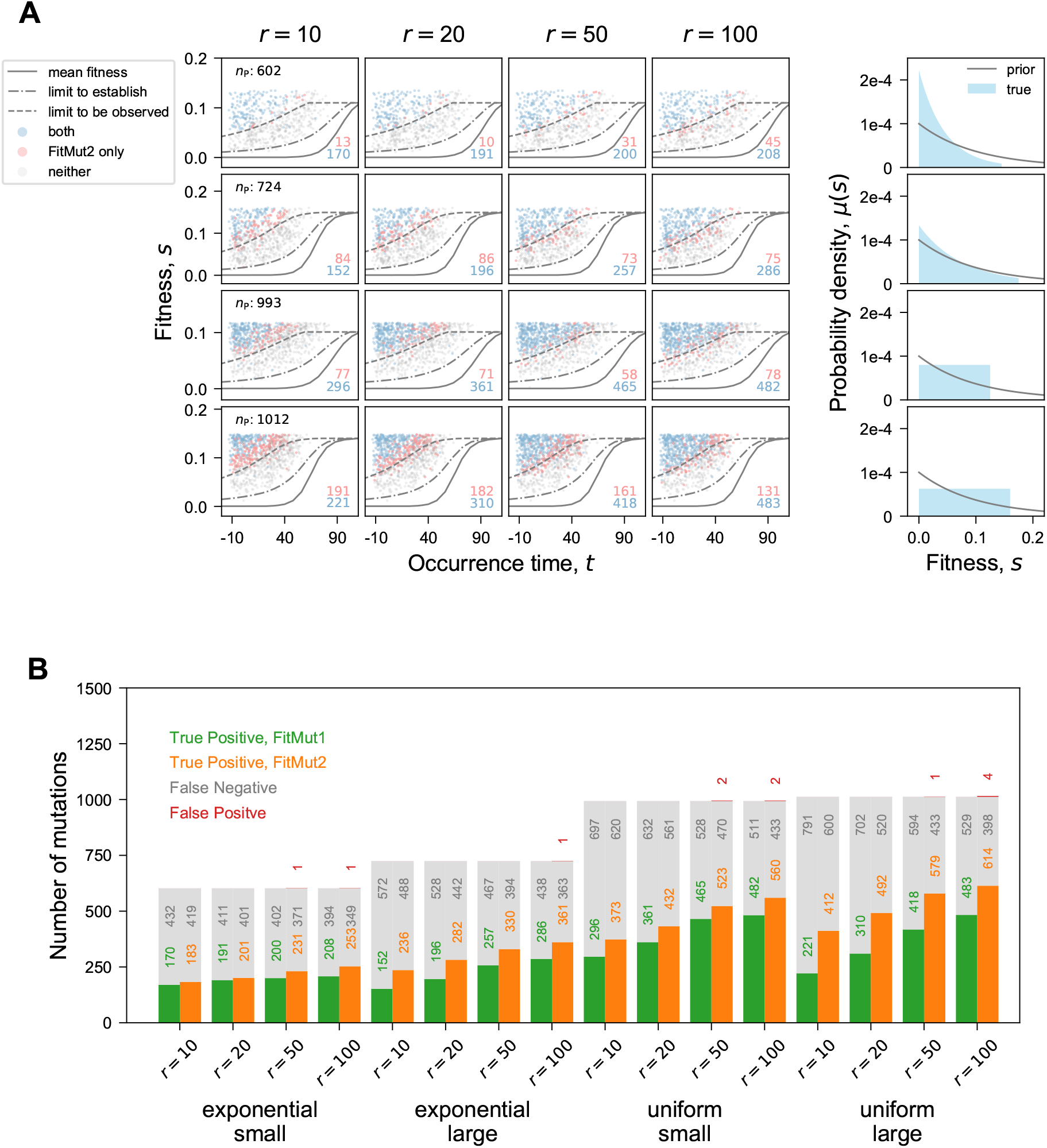
Detection ability for identifying adaptive mutations. (A) Each panel in the 4 × 4 array corresponds to one simulation (Methods:B). Each point represents an adaptive mutation that occurred and established in the simulation. Points are colored according to whether they were identified by both methods (blue), only by `FitMut2` (pink), or by neither (grey) (no point that only by `FitMut1`); their counts are shown in the right bottom corner of each panel. *n_P_* represents the total number of established mutations for a given DFE. The three lines indicate the mean fitness (solid, 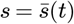), the boundary above which mutations must occur in order to establish (dot-dashed, 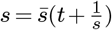) and the boundary to be observed (short-dashed, 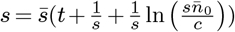). The 5th column shows *μ*(*s*) and the prior 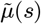 for each row. (B) Direct comparison of the detection ability between both algorithms.

We also compared the number of adaptive mutations missed by each algorithm, as well as the number of false positives (Figure 5B). It should be emphasized that the vast majority of false negatives fall below the observation limit as detailed below, and are not expected to be detected. Additionally our algorithm reports an estimate of the uncertainty in our inferred values of *s* and *τ*, which is based on the Hessian of the posterior likelihood at the optimal values of *s* and *τ* (Section S6).

### D. FitMut2 identifies mutations closer to the limit of detection

The clonal interference regime imposes a limit on the fitness effects that can establish, as well as those that can be detected. Although mutations occur throughout the experiment, only those that rise to a large enough frequency can be detected. Following (Levy et al., 2015), we restate the rough requirements for establishment and detection of a beneficial mutant. For an adaptive mutation with fitness effect *s* in a birth-death process with individual offspring-number variance per generation 2*c*, it takes 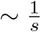 generations for the mutation to establish, and another 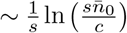 generations for the established mutation to sweep through an appreciable fraction of the lineage to be detectable. Here, 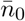 is the average effective lineage size given in our case by 100. Therefore, typically, a mutation should satisfy 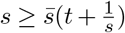 in order to establish, and also satisfy 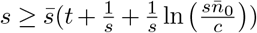 in order to be observed. Paired values *s* and *t* that satisfy these equations are found numerically and shown in Figure 5A. We see that in Figure 5A, as expected, identified adaptive mutations lie largely above the highest line.

`FitMut2` is better able to identify mutations closer to the bound of detectability, as evidenced by the predominance of red points just above the uppermost curve in Figure 5A. However, it has a nonzero false positive rate when the read depth is large (Figure 5B). One way to combat the identification of false positives is to increase the threshold for designating a mutation as adaptive. In this work, the threshold is 0.5: we call a lineage adaptive if our Bayesian estimate says that the probability of it being adaptive is greater than 0.5. Increasing this threshold should lower our false positive rate.

## Discussion

In this work we have extended a previously-devised algorithm to infer mutation effects and establishment times from lineage trajectories over time. Using simulated data we have shown that our new algorithm `FitMut2` performs better than the previous version `FitMut1` when the read depth is low or the distribution of fitness effects is broad. By inferring the population mean fitness and single mutation effects self-consistently, instead of relying on the decline of neutral lineages, we can apply our algorithm to datasets with shallower sequencing, rapidly adapting populations, or smaller initial lineage sizes. However there are a number of aspects of this algorithm that deserve additional comments.

### Branching process model

The model of a growing lineage described in S10 that we use to derive the distribution of read number conditional on a past measurement assumes that the lineage is reproducing and dying at constant rates in time, and that the difference between these rates constitutes the fitness. However, in the serial batch culture experiment (and in simulation), the population grows by two orders of magnitude (~ 2^8^) with minimal death every batch culture cycle — and this changes theoretical expectations for the distribution of offspring number from one measurement to the next. As discussed in S9, this can mostly be absorbed into an effective population size which is *g* times the bottleneck size. However for large-effect mutations, the growth stochasticity during a single cycle may obey different statistics and a more careful analysis of the effective parameters is needed.

### Independence of Sequencing Noise Across Time Points

One shortcoming of our approach (and that of `FitMut1`) is the assumption that the distribution of read number for a given lineage at a time point *t_k_* depends only on the number of reads counted at the previous time point: this approximation makes the problem of maximizing likelihood much more tractable. However, sequencing noise at different time points is uncorrelated: therefore if sequencing noise caused the read number to be large at the previous time point, there is no reason to believe that the subsequent measured read number would have a larger mean. The independence of sequencing noise across time points could be used to our advantage, allowing us to separate the stochasticity from cell division (which is of biological interest) from that due to sequencing noise. Though we have not pursued this direction in the current work, it remains a promising avenue. Previous work (Levy et al., 2015) has measured the value of *c* through doing multiple sequencing replicates — but the presence of multiple time points could allow us to circumvent the need for this extra sequencing.

### Choice of prior

To infer the fitness effect and establishment time of a mutation, we must choose a prior distribution 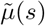 on which to run our Bayesian inference. In this work we consistently used an exponential prior 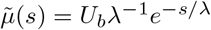 with *λ* = 0.1 and *U_b_* = 10^-5^. The prior for *s* and *τ* was then 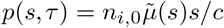 with *s/c* factoring in the establishment probability and *n*_*i*,0_ accounting for effective size of lineage *i* at *t*_0_. The prior does not depend on *τ*, whose prior distribution we took to be uniform between −100 and 112 generations.

The use of an exponential prior assumes that there are no very large mutations — because if there were, we would be increasingly unlikely to recognize them. Therefore in situations where the distribution of fitness effects is broader, an exponential prior may fail to identify many adaptive mutations, and a uniform prior may be more appropriate. The effect of the prior is further discussed in Section (S7), where we conclude that our choice of prior makes little difference for the inferred *s_i_* in the majority of adaptive lineages.

To lessen the arbitrariness of our choice for 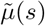, we could consider having a dynamically updated prior that starts as uniform and is updated based on the mutations identified as adaptive over successive iterations of our algorithm. It is conceivable that this would further increase our detection power. But there would be issues of low resolution in this empirically determined prior distribution, particularly for early iterations. Future work is needed to investigate how to iteratively update the prior distribution, and whether this further improves estimation accuracy.

### Rebarcoding and measurements of epistasis

As mentioned previously, our simulations allowed a maximum of one beneficial mutation per individual, since the DFE for a second mutation might be substantially different from that of the first (Aggeli et al., 2021). The effects of recurrent beneficial mutations can be studied systematically using genetic re-barcoding of lineages (Nguyen Ba et al., 2019), where a similar fitness-estimation algorithm has been implemented by iteratively inferring mean fitness while also identifying lineages with a beneficial mutant. However, this algorithm is used as a rough heuristic to identify adaptive lineages rather than to infer the fitness effects and establishment times of individual mutations, and so our method complements this approach in cases where the values of the inferred parameters are desired with higher accuracy.

### Computational performance

The most computationally expensive step in both `FitMut2` and `FitMut1` is the evaluation of the probability of being adaptive for each lineage, and the subsequent maximization of the posterior likelihood for those deemed adaptive. However, this step can readily be parallellized (in both `FitMut2` and `FitMut1`) since each lineage may be handled independently. This substantially speeds up our algorithm, and we have included an option to parallellize computation using the python package `multiprocess`, which distributes iterations of the longest for loop in the program over multiple CPUs if available. With parallelization enabled, on our simulated dataset of 4 × 10^4^ lineages sampled over 15 time points, `FitMut2` took around 1 minute per iteration on a laptop with 8GB of memory. In comparison, `FitMut1` took around 15 minutes in total.

## CODE AND DATA AVAILABILITY

All of our code for simulations and inference, as well as the code to generate the figures in this paper, is available at the github website https://github.com/FangfeiLi05/FitMut2.

## ACKNOWLEDGEMENTS

This work was supported by NIH R01 AI136992 to GS, and by NSF PHY-1607606. Some of the computing for this project was performed on the Stanford Research Computing Sherlock cluster. We thank Daniel S Fisher for useful discussions and suggestions.

## Supplementary Information

### S1. Definition of Notation

**Table S1.** Definitions of commonly used quantities.

| Symbol | Definition |
| --- | --- |
| $T$ | total number of generations in evolution experiment |
| $L$ | number of barcoded lineages |
| $g$ | number of generations between successive cell transfers |
| $t_0, t_1, t_2, \dots, t_K$ | list of sequencing time points ($K + 1$ time points in total) |
| $r_{i,k}$ | read number of lineage $i$ at $t_k$ |
| $R_k$ | total sequencing read depth at $t_k$, given by $R_k = \sum_{i=1}^L r_{i,k}$ |
| $B_k$ | total number of cells transferred at bottleneck occurring at time $t_k$ |
| $N_k$ | effective population size at time $t_k$, given by $N_k = gB_k$ |
| $f_{i,k}$ | frequency of lineage $i$ at $t_k$, given by $f_{i,k} = n_{i,k}/N_k$ |
| $n_{i,k}$ | effective lineage size of lineage $i$ at time $t_k$ |
| $s_i$ | fitness effect of a mutation arising in lineage $i$ |
| $\tau_i$ | establishment time of a mutation in lineage $i$ |
| $\bar{s}(t)$ | mean fitness of the population at time $t$ |
| $\mu(s)$ | distribution of fitness effects (DFE) of mutations |
| $U_b$ | total beneficial mutation rate, given by $U_b = \int_0^\infty \mu(s) ds$ |
| $2c$ | single individual offspring number variance per generation in the branching process model |
| $2\kappa_k$ | phenomenological parameter capturing per-read variance in offspring number<br>between times $t_{k-1}$ and $t_k$ |
| $\mathcal{E}_u^v$ | reduction in frequency due to mean fitness, defined as $\mathcal{E}_u^v = \exp \left[ - \int_u^v \bar{s}(\eta) d\eta \right]$ |
| $\Theta_i$ | indicator variable for the establishment of a mutation in lineage $i$ |

### S2. Definition of Fitness

Evolutionary theory commonly uses two different definitions of fitness: *Malthusian* fitness and *Wrightian* fitness. Let *t* denote elapsed time (measured in generations), and *n*(*t*) denote the number of cells at time *t*. Malthusian fitness *s_m_* is defined as the exponential growth rate of the population, and Wrightian fitness *s_w_* is defined as the number of offspring of a cell per generation. So for deterministic growth without death these definitions imply

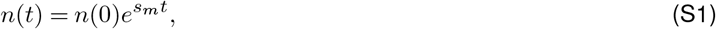

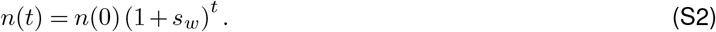

The fitness of a mutation in our simulations is defined in the Wrightian sense, which is natural for discrete generations. The branching process model we employ uses fitness in the Malthusian sense, which arises naturally in continuous time. Malthusian and Wrightian fitness are related by *s_m_* = ln(1 + *s_w_*) and are approximately equal when the fitness effect per generation is small, as is the case in our analyses. For fair comparison, we convert the mean fitness and fitness effect in the simulation from Wrightian fitness to Malthusian fitness when comparing the ground truth of the simulation to the inferred value.

In a well-mixed environment without ecological interactions, all cells in the population compete against the mean fitness. For an adaptive lineage that originated from a single mutant cell, when the fitness of lineage *i, s_i_*, is larger than the mean fitness, the lineage increases in frequency. Once the mean fitness passes the fitness of the lineage, the lineage begins to decrease in frequency. Over time, the mean fitness increases as fitter adaptive lineages become enriched. If the total population size is fixed and we track lineage frequencies {*f_i,k_*} at each time point subject to ∑_*i*_ *f_i,k_* = 1, one can see that they obey the equation

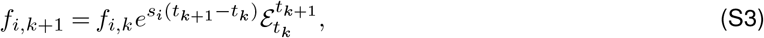

where 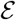 is defined in Table S1 and 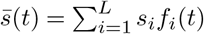, which varies continuously in time. Equation S3 is used to estimate the average read number of a lineage conditioned on its read number at the previous time point — however since we only infer the mean fitness at time points that are sequenced, we interpolate 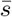 linearly between these points of reference. Under this approximation,

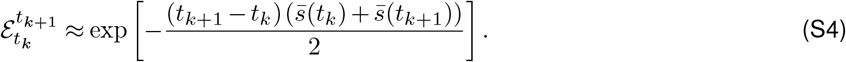

### S3. Effective Process

As discussed in the SI of (Levy et al., 2015), the process of population growth for *g* generations followed by 1 : 2^*g*^ dilution can be described as a branching process with the correct effective values of *n*(*t*) and 2*c* (Section S10). The effective lineage size *n*(*t*) is *g* times the number of cells transferred at the dilution, and the offspring number variance per individual per generation, 2*c*, is given by the per-individual variance *per growth cycle* from the batch culture. Equivalently, one could define the effective size of each lineage as its size at bottleneck, and 2*c* reduced from the variance per cycle by a factor of *g*. In this work, we use the former convention.

In our simulated model with stochastic cell doubling and dilution noise, the per-individual variance in offspring number per cycle can be calculated using the law of total variance, and is 2*c* ≈ 2, with *g* generations of stochastic cell doubling introducing a variance of ≈ 1 for *g* large, and 1 : 2^*g*^ dilution introducing another variance of 1. Therefore if the branching process model is to describe the simulated process, we should set *c* = 1. However in real data the value of *c* is unknown, and is of biological interest. Inferring it from our data is a future goal as mentioned in the Discussion. In the current framework, setting *c* =1 only affects the establishment size of a beneficial mutation during the inference process, and we therefore expect that it does not much change our results.

### S4. Parameterizing Noise

Barcoded evolution experiments involve two processes: 1) a series of batch cultures, where in each cycle the population grows for about *g* generations and then is diluted by a factor of 2^*g*^ into the next batch, and 2) sampling and preparation of cells via DNA extraction, PCR and sequencing. As discussed in the supplementary information of (Levy et al., 2015), both processes introduce stochasticity in the number of reads per barcode at each time point. Dealing with this stochasticity is essential for detecting adaptive mutations and estimating their mutational parameters. Here we follow the approach of `FitMut1`, where we assume that the variance of *r_i,k_* is proportional to 〈*r_i,k_*〉, its expected value. This relationship is consistent with birth-death stochasticity arising from a branching process, and also with the noisy doubling of DNA during PCR. The constant of proportionality is given by a phenomenological parameter 2*κ_k_*, which captures all sources of noise (cell growth and dilution, DNA extraction, PCR and sequencing). Under this noise model, Var(*r_i,k_*) = 2*κ_k_*(*r_i,k_*) for every lineage *i*. In order to determine the values of the *κ_k_* for each time point *t_k_*, we first find all lineages with read number *r*_*i,k*–1_ = *q* at *t*_*k*–1_. Then we estimate the mean *μ* and variance *σ*^2^ of *r_i,k_* conditioned on *r*_*i,k*–1_ = *q*. Then, conditional on *q*, *κ_k_* = *σ*^2^/(2*μ*). We average *κ_k_* over all *q* ∈ [20,30], which are likely to correspond to neutral lineages for typical read depths. Although we only determine the *κ_k_* once and for all at the beginning of our inference algorithm, we have found that our results are quite insensitive to the values of the {*κ_k_*} inferred.

### S5. Algorithm Details

Here we describe in detail the algorithm used by `FitMut2`. The basic premise is Bayes’ theorem: we wish to know the probability that each lineage was adaptive, conditional on our observed lineage trajectory {*r_i,k_*}. In this and the subsequent section, we assume that the lineage index is always *i*, and suppress the *i* subscript. Bayes’ theorem tells us that for a lineage with observed lineage trajectory {*r_k_*},

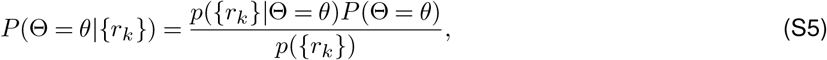

where Θ is an indicator variable that is 1 if the lineage contains an established mutation and 0 otherwise.

When *θ* = 1, we can further decompose this adaptive hypothesis into an integral over all possible ways to be adaptive, with a prior distribution *p*(*s,τ*) over *s* and *τ*. Thus,

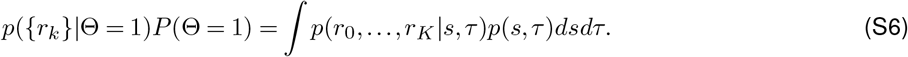

When *θ* = 0 we use the prior *P*(Θ = 0) ≈ 1, since the vast majority of lineages do not obtain a beneficial mutation (or equivalently ∫*p*(*s,τ*)*dsdτ* ≪ 1). Therefore, we reduce the problem of inference to calculating 1) the probability of the *{r_k_}* given the neutral hypotheses and 2) the probability of the *{r_k_}* given the adaptive hypothesis with a mutation having fitness effect *s* and establishment time *τ*. Together with a prior distribution over *s* and *τ*, this gives us the tools to assess the probability that a lineage is adaptive. We next explain how to calculate these terms.

#### Neutral hypothesis

To calculate *p*({*r_k_*}|Θ = 0), we make an important approximation. First we note that

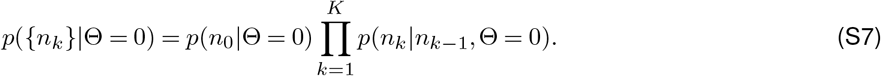

In other words, the *cell number* at each time point obeys a Markov process, and the distribution of *n_k_* only depends on the value of *n*_*k*–1_. Note that strictly speaking, the analogous statement for the read numbers {*r_k_*} is false: the distribution of *r_k_* does not only depend on *r*_*k*–1_, due to independent sequencing noise introduced at each time point.

Nonetheless, we make the approximation that the joint distribution of the {*r_k_*} factorizes in the same way as that of the {*n_k_*}. Though we expect that this assumption become less accurate a low read depth, empirically our algorithm still seems to be accurate over the range of parameters we have tested. We additionally investigated an alternative method that does not make this assumption, and instead tries to find the sequence of cell numbers {*n_k_}* that is most likely to have produced our data. However, we have not pursued this method in the current work, due to its infeasibility under the adaptive hypothesis. Instead, we define the likelihood of our data given the neutral hypothesis as

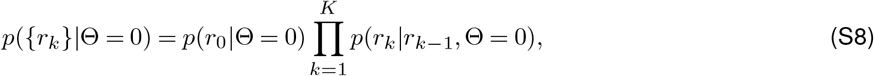

where, according to Equation S28,

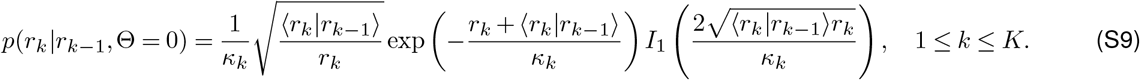

Here *I*_1_ is a modified Bessel function of the first kind, and the notation 〈*r_k_*|*r*_*k*–1_〉 indicates the expected value of *r_k_* conditional on the value of *r*_*k*–1_ and Θ = 0. This value is given by the expression for the decline of a neutral lineage in the presence of an increasing mean fitness:

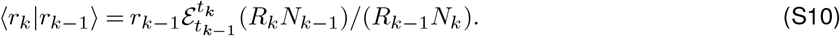

We can then take the product of these conditional distributions as specified in Equation S8, in order to get the likelihood of our data under the neutral hypothesis. However we must deal separately with the first time point which gives us a factor of *p*(*r*_0_). This is discussed below.

#### Adaptive hypothesis

For the adaptive hypothesis we have

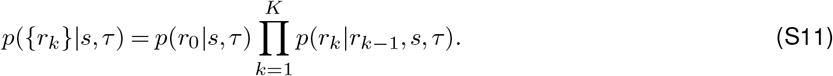

When an adaptive mutation occurs and establishes in a lineage, that lineage can contain both mutant and neutral cells which are indistinguishable on the basis of barcode alone. Before the establishment, the neutral cells comprise almost the entire lineage, while after an adaptive mutation establishes, the mutant cells begin to sweep through the lineage.

In order to calculate the likelihood of our data under the adaptive hypothesis, we use Equation S9, conditioned on a value of *s* and *τ* rather than Θ = 0, and with the conditional expectations 〈*r_k_*|*r*_*k–1*_) calculated under the assumption of a given *s* and *τ* (Equation S12). Assuming that an adaptive mutation with fitness effect *s* establishes at time *τ* in a lineage, the average read number of a lineage at time point *t_k_* is

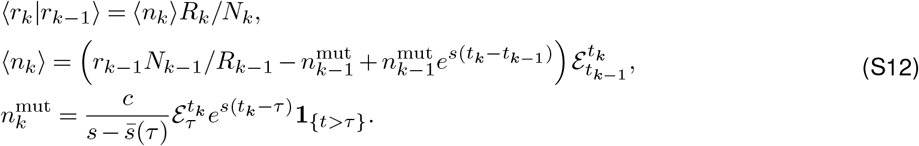

Here, **1**_{*t*>*τ*}_ is an indicator variable that is 1 if *t* > *τ* and 0 otherwise, and we have used the fact that the establishment size of a lineage beyond which it grows deterministically is given by 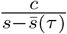.

Once 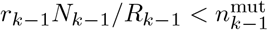, we assume that 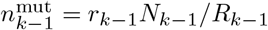. Furthermore, we impose a lower cutoff on 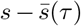 to avoid very large establishment sizes.

Since we also want to calculate likelihoods for *τ* < 0 (corresponding to mutations in the pre-growth phase), we need the mean fitness before *t* = 0 when calculating 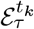: we assume that the mean fitness is 0 before the experiment begins.

As in the case of the neutral hypothesis, we must deal with the first sequencing time point *t*_0_, for which we do not have a previous time point on which to condition. Therefore for the first time point we define

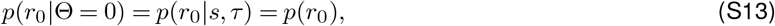

with

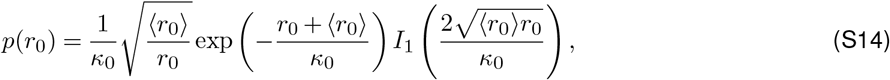

and 〈*r*_0_〉 = *r*_0_: in other words the mean *r*_0_ is approximated by the measured value of *r*_0_ for each lineage. *κ*_0_ is assumed to be 2.5, which is empirically a typical value. Estimating the most likely value of *n*_0_ and the resulting distribution of *r*_0_ from the data is an interesting problem which we leave for future work.

Therefore under the adaptive hypothesis, the probability of observing our data is given by

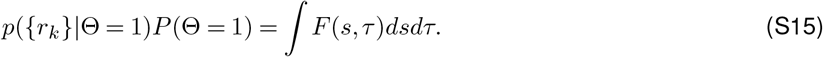

where we define the *posterior likelihood F*(*s,τ*) as

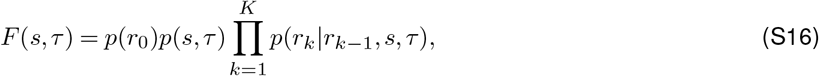

with prior distribution

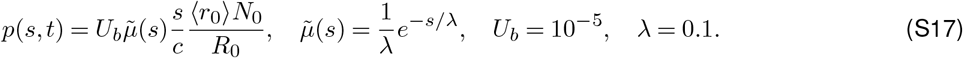

Therefore to calculate the probability that a lineage is adaptive, we numerically integrate *F*(*s,τ*) for that lineage over *s* and *τ*. Since this calculation for each lineage is independent, it can be readily parallelized as discussed in Discussion.

The two terms *P*({*r_k_*}|Θ = 0)*P*(Θ = 0) and *P*({*r_k_*}|Θ = 1)*P*(Θ = 1) must sum up to the marginal probability *P*(*{r_k_*}), which then allows us (using Equation S5) to calculate the *probability* that our lineage is adaptive or neutral given the data, that is, *P*(Θ = 0|{*r_k_*}) and *P*(Θ = 1 |{*r_k_*}). If *P*(Θ = 1 |{*r_k_*}) *>* 0.5, we claim that the lineage contains an established mutation.

The fitness effect and establishment time that we infer for this lineage are given by the *s* and *τ* that maximize the posterior likelihood *F*(*s,τ*). This maximization is discussed in the next section.

#### Numerical Optimization

The fitness effects and establishments times of adaptive mutations are inferred by optimizing the posterior likelihood (S5) using the Nelder-Mead algorithm, which finds the maximum of a function of *n* variables by constructing a simplex of *n* + 1 points and then updating the vertices of the simplex according to a set of prescribed rules. (Nelder and Mead, 1965). Though this algorithm is not guaranteed to find the global optimum, it is widely used for nonconvex optimization and finds optima without taking derivatives of a landscape. In some of our work we observed sensitivity of the maximization algorithm to the simplex with which it was initialized, but in our current code we have chosen the initial conditions to robustly yield actual optima.

Figure S5 shows the landscape of the log-posterior likelihood for an adaptive lineage in which an adaptive mutation established, as well as the landscape of the log-posterior likelihood for a neutral lineage. We see that *F_i_*(*s,τ*) is not particularly sharply peaked in all directions — in fact its density is strongly concentrated along a one-dimensional curve in two dimensions. Though the algorithm accurately finds the maximum in the likelihood landscape, there is some mismatch between the location of this maximum and the true values of the fitness and occurrence times.

#### Updating the Mean Fitness

Since the algorithm proceeds iteratively, we need a rule for updating the mean fitness after the fitnesses of individual lineages are inferred. Specifically, at the end of each iteration, we estimate the effective number of mutant cells for each lineage 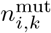 for each time point *t_k_* given the *s_i_* and *τ_i_* that were inferred. Then we update the mean fitness as

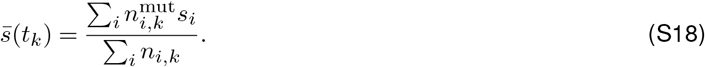

### S6. Error Estimation

In addition to estimating *s* and *τ* for each adaptive lineage, we provide uncertainties of our estimates, which can be calculated via the Hessian matrix of ln*F*, evaluated at maximal point (*s*_0_, *τ*_0_) found by our optimization algorithm. The Hessian is given by

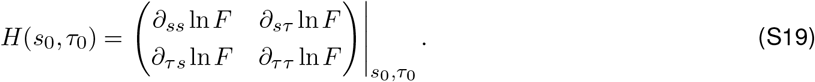

The eigenvalues of *H* give information about how sharp of a maximum in ln*F* we have found — if the maximum is very shallow we are more uncertain of its location, and therefore of our estimates of *s* and *τ*. Following (Levy et al., 2015), we calculate the eigenvectors 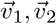 and eigenvalues *λ*_1_, *λ*_2_ of *H*. By projecting the eigenvectors along the *s* and *τ* directions of our parameter space and using the fact that *F* is locally Gaussian with scale 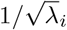 in the direction 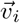, we define the error in *s* to be 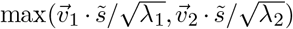 and the error in *τ* to be 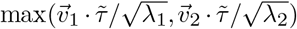 where 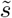 and 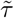 are the unit vectors (1,0) and (0,1) respectively.

### S7. Effect of Prior

The choice of prior 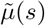 affects the landscape over which we optimize ln*F*(*s,τ*). Ignoring logarithmic terms, which have a negligible effect on the location of the maximum of *F*, we can expand our objective function as

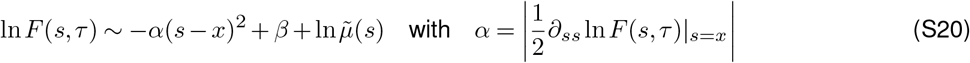

in the neighborhood of the optimum, where *x* is the optimal *s* that would be found under a uniform prior. Therefore if we use a prior 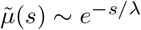, we find that the optimal *s* where *F* is extremized shifts by an amount 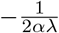. If the lineage does not contribute much to the mean fitness, this shift in the fitness of lineage does not affect the optimization landscape and the prior that we have chosen simply shifts 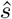 by an amount that depends inversely on *a* and *λ*.

From Figure S5 we estimate that *α* ≈ 20/.05^2^ = 8000. Therefore varying *λ* from 0.1 (as in this work) within an order of magnitude would have little effect on 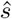.

### S8. DFE estimation

Let *f*(*s,t*) be the total fraction of cells in the population with fitness effect in the range [*s,s* + *δs*] at the time *t*. According to (Levy et al., 2015),

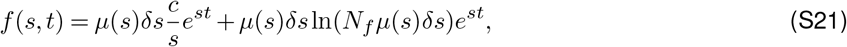

where the first term on the right side accounts for mutations during the evolution, and the second term accounts for mutations that arose during pregrowth. *N_f_* = 10^12^ is the maximum population size after barcoding, before the separation of the replicates.

Thus, the DFE *μ*(*s*) satisfies the

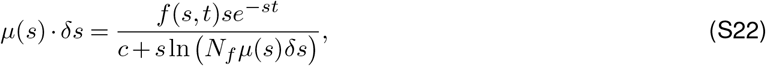

which can be solved numerically to yield *μ*(*s*). Here, Equation S22 is an approximation that without considering the effect of the mean fitness. Therefore, to infer *μ*(*s*) from our results, we choose the time point *t* = 32, at which time the mean fitness is still small.

### S9. Simulation Details

In our simulations, during 16 generations of pregrowth, the number of offspring of a single cell with fitness *s* is distributed as Pois(2(1 + *s*)). This allows lineage size to fluctuate, and introduces variability in the lineage size going into the pooled batch culture. At the end of the pregrowth phase, each cell *i* has grown into a colony of size 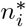. To initialize the evolution experiment, an average of 100 cells per barcode are sampled: *n_i_*(0) cells from each lineage are sampled with

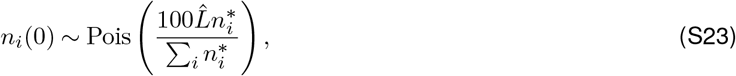

where 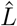 is the number of non-extinct colonies. 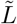 is then defined as the number of non-extinct lineages with *n_i_*(0) > 0. During each batch culture cycle, growth noise is simulated by updating the number of descendants of a single cell according to

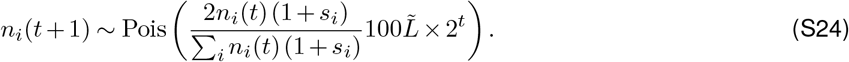

After *g* generations, the cells which get transferred to the next batch are sampled randomly from the final population and are Poisson distributed with mean *n_i_*(*g*)/2^*g*^.

Of the four DFEs in our simulation, two are truncated exponential distributions: *μ*(*s*) ~ exp(–*s*/0.045) with *s* ∈ (0,0.145), and *μ*(*s*) ~exp(–*s*/0.075) with *s* ∈ [0,0.175]. The othertwo are uniform between on intervals (0,0.125) and (0,0.16) respectively. All DFEs have 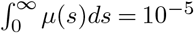.

In order to determine whether a mutation has established or not in simulation, we compare its instantaneous effective cell number *n*(*t*) to 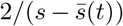, and if it crosses this size we record it as established — this heuristic means that its probability of extinction would be about *e*^−2^ if 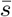 stopped increasing thereafter (Methods:B). When we infer the mutational identity of lineages from experimental data, the establishment size used in our inference algorithm is 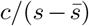. However, for real data we do not know *a priori* what the value of *c* is in experimental data, and our algorithm assumes that *c* = 1.

### S10. Distribution of offspring from a single individual

The time-homogeneous birth-death process is a simple model to understand the growth of a lineage that is subject to both stochastic drift and systematic selection. Let *n*(*t*) denote a random variable that represents the number of individuals at the time *t* (where time is now taken to be continuous and in units of generations). Assume that each individual has a birth rate *k_b_* and death rate *k_d_*, and individuals grow or die independently of one another. In a small interval of time *δt* ≪ 1/*k_b_*, 1/*k_d_*, each individual divides with probability *k_b_δt*, dies with probability *k_d_δt*, and does nothing with probability 1 – (*k_b_* + *k_d_*)*δt*. Let *B_j_* and *D_j_* respectively denote Bernoulli random variables indicating birth or death in the interval [*t,t* + *δt*] for the *j*th individual. As done in (Levy et al., 2015), we define a moment generating function of this birth-death process

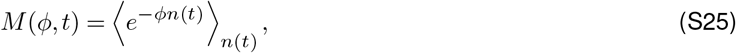

with 〈*X*〈_*Y*_ denoting an average of the random variable *X* over the random variable *Y*. Since our branching process obeys the equation 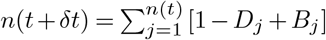, we derive a partial differential equation for this moment generating function

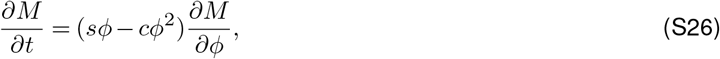

where *s* = *k_b_* – *k_d_* and *c* = (*k_b_* + *k_d_*)/2. We then solve this equation by the method of characteristics

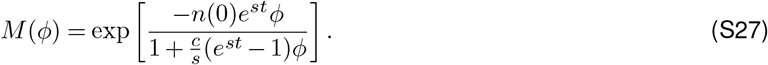

We finally get probability distribution *p*(*n*) of the cell number *n* at time *t*, starting from *n*(0) at time 0, is given by the inverse Laplace transform of *M*(*φ*),

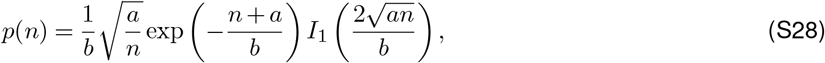

where *a* = *n*(0)*e^st^*, 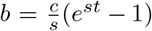. Here, *I*_1_(*x*) denotes a modified Bessel function of the first kind 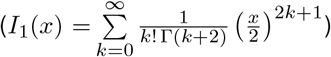. The mean of this distribution is *a* and its variance is 2*ab*. For *n* large compared to *b*^2^/*a*, our distribution takes the limiting form

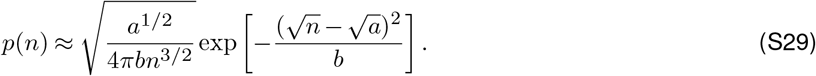

Equation S29 is the theoretical distribution that `FitMut1` used for the number of reads at the current time point *t_k_* conditioned on the previous time point *t*_*k*–1_, where *a* is the mean number of reads at the current time point, and *b* is replaced by the *κ_k_*. Equations (S10) and (S12) are used to calculate the expected number of reads corresponding to a particular barcode at the current time point, conditional on the read number at the previous time point. In `FitMut2`, we use Equation S28, which is more accurate for small read numbers.

Our branching process can also tell us what the establishment size of a lineage is, above which it is destined to fix. Note that *P*(*n*(*t*) = 0) is the extinction probability by time *t*. We can calculate this probability directly from the moment generating function according to

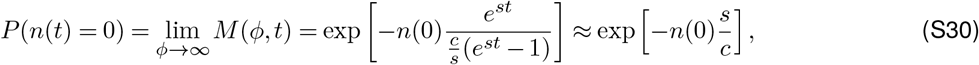

where the last approximation obtains when *t* ≫ 1/*s*, which corresponds to eventual probability of fixation. Therefore the lineage of a single mutant cell (carrying an adaptive mutation with fitness effect *s* ≪ *c*) goes extinct with probability ≈ 1 – *s/c* and establishes with probability *s/c*. Furthermore, once a lineage reaches size *c/s*, it is unlikely to go extinct. If we define an establishment size *c/s* then the establishment time *τ* is defined through the relation 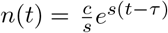 (with *n*(0) = 1 meaning that the mutation occurred at time 0), which yields a *t*-independent distribution for *τ* – *t* as *t* → ∞ (Desai and Fisher, 2007; Levy et al., 2015). *τ* – *t* is asymmetrically distributed around 0 with width scaling as 1*/s*, and *τ* – *t* much more likely to be large and positive than large and negative on this scale. However the most likely value of *τ* – *t* is 0, which allows us to compare the occurrence time from simulation to establishment time from inference.

## Supplementary Figures

**Figure S1.**
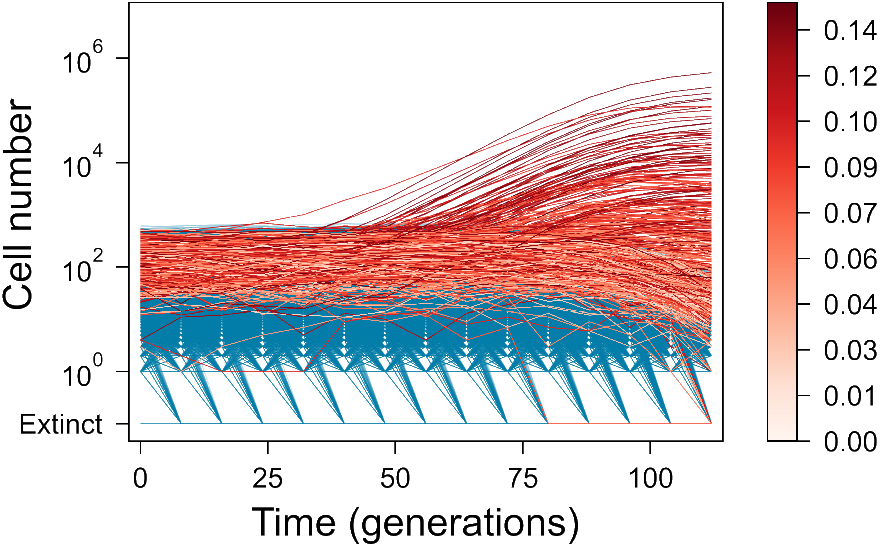
Trajectories of lineages. Lineages that contain an adaptive mutation that established are colored by the fitness effect of the adaptive mutation in the lineage. Neutral lineages are shown in blue.

**Figure S2.**
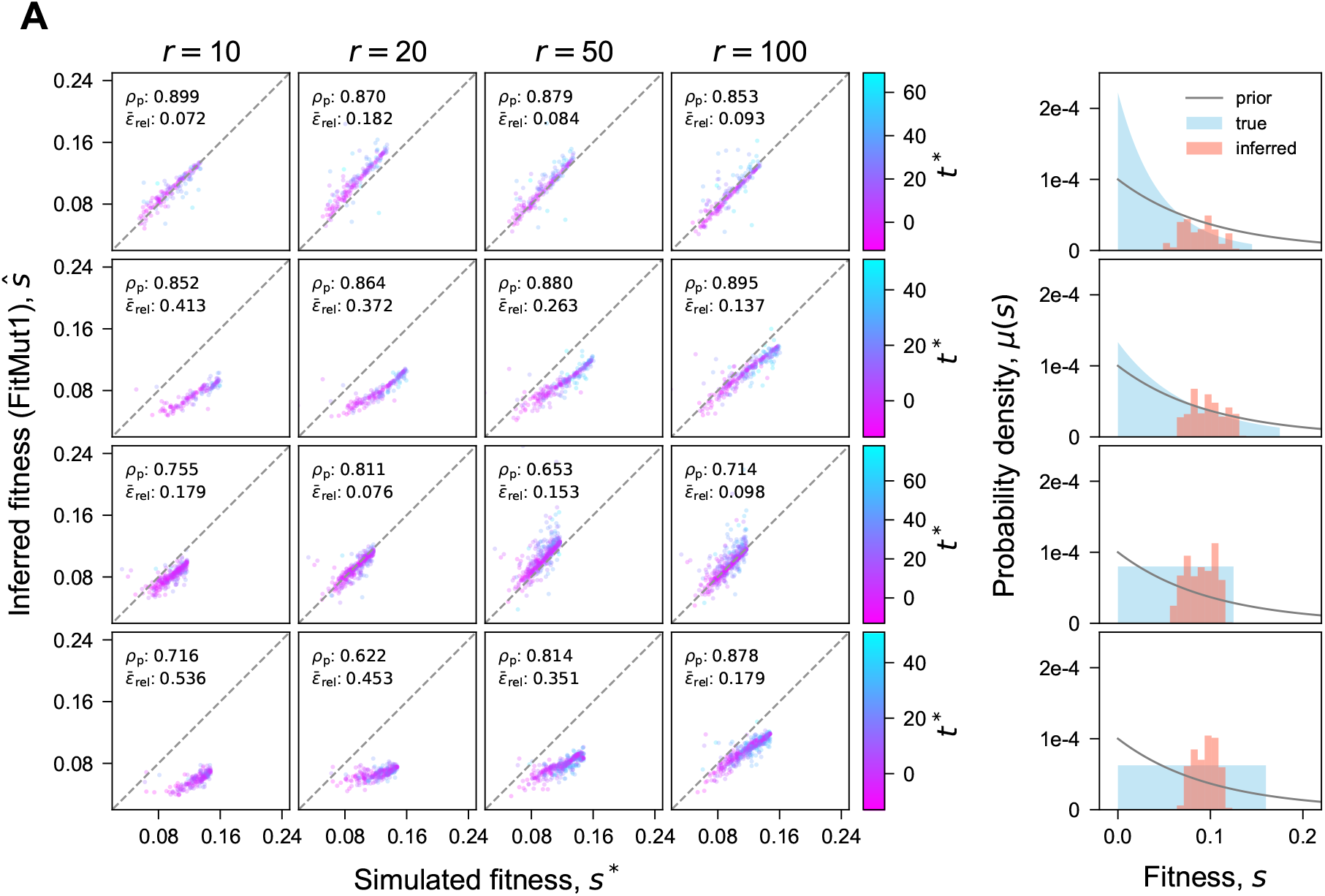
Inference accuracy of (A) fitness effects and (B) establishment times (FitMut1). Comparison of inferred *s* and *τ* with simulation. Compare with equivalent Figure 3 for `FitMut2`. See caption of Figure 3 for parameter definitions. The agreement between true and inferred values is significantly worse than the same results using `FitMut2` (Figure 3).

**Figure S3.**
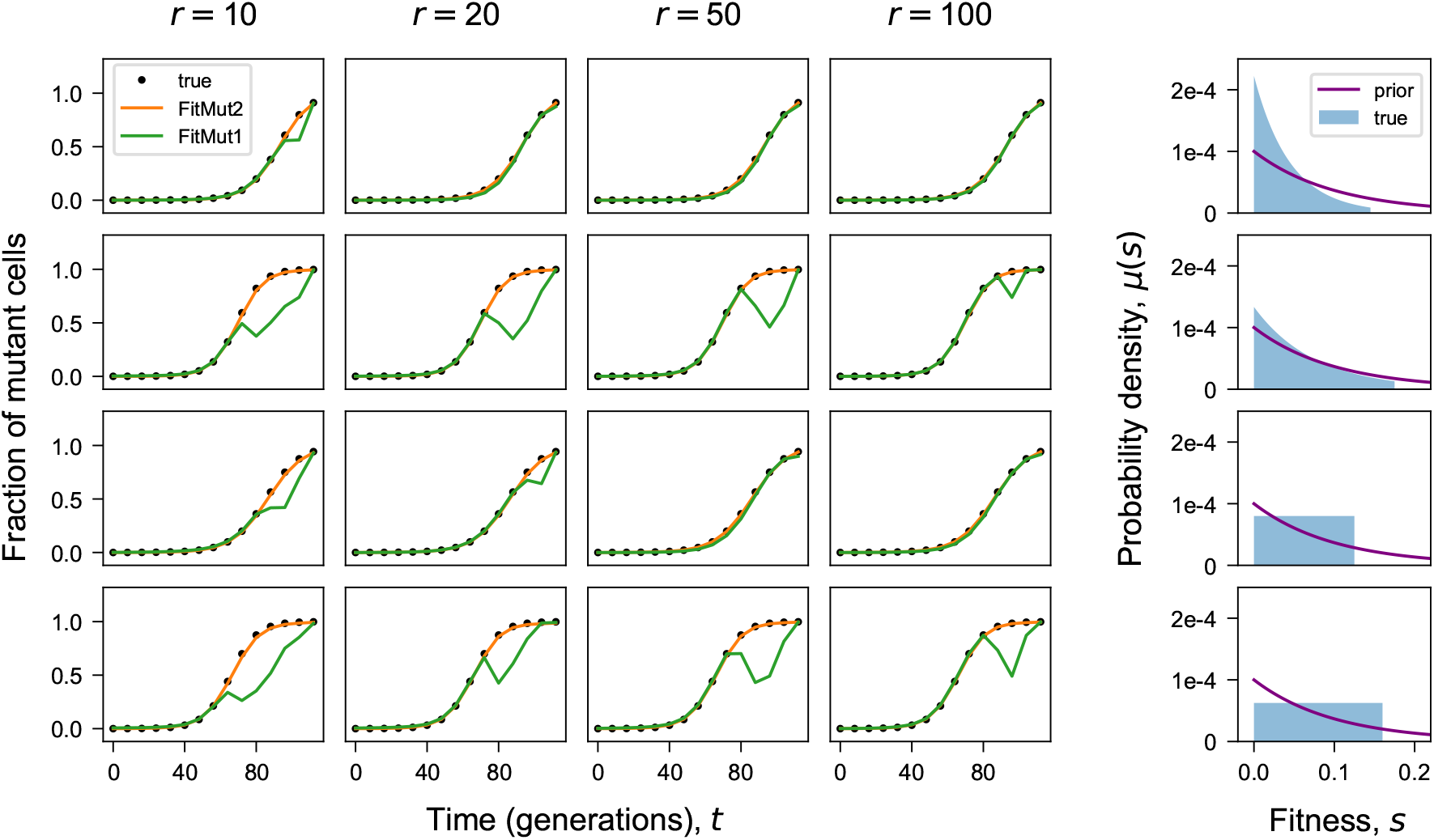
Mutant fraction of the population. Comparison of the estimated and true total frequency of mutant cells in the population over time. Each panel in the 4 × 4 array corresponds to one simulation (Methods:B). The 5th column shows the DFE *μ*(*s*) used in each simulation condition, and the prior 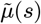 used for both `FitMut2` and `FitMut1`.

**Figure S4.**
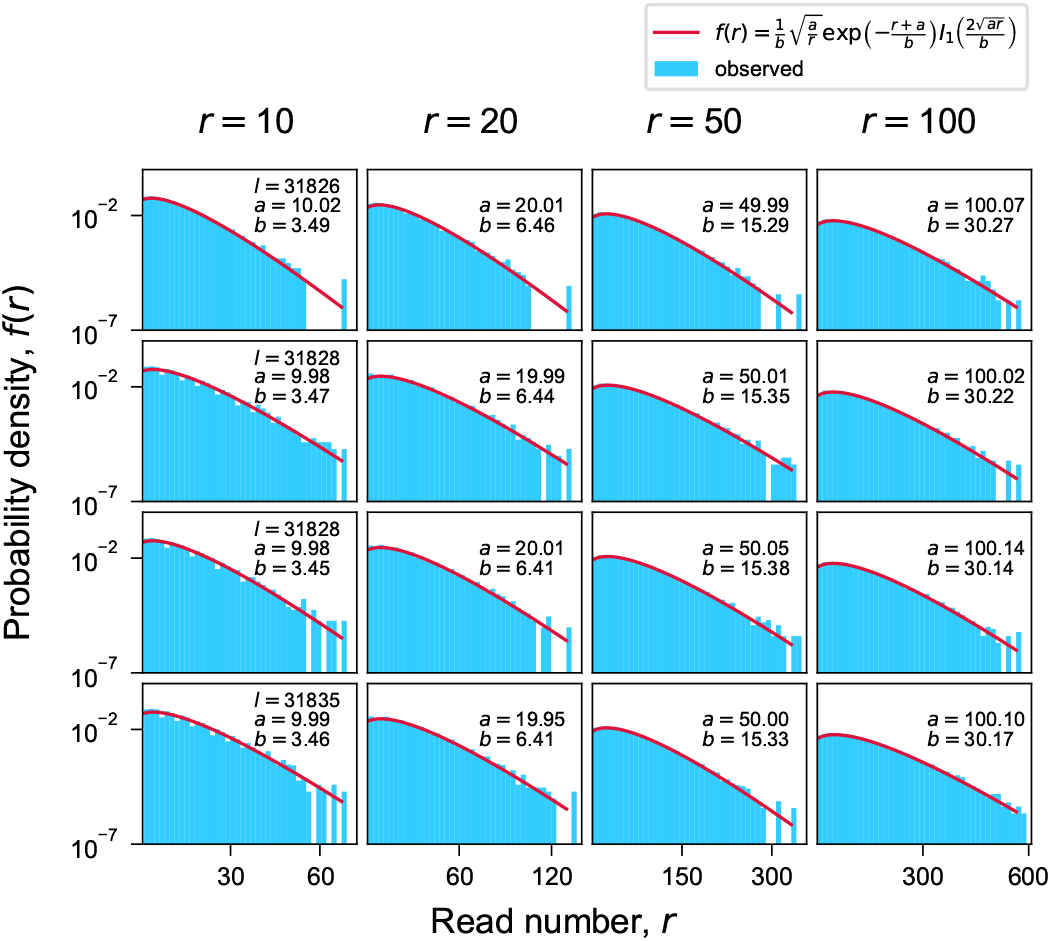
Probability distribution of read number. Probability distribution of the read number *r* at *t* = 0. Each panel in the 4 × 4 array corresponds to one simulation (Methods:B). The blue histogram is a plot of simulated data. The theoretical probability distribution (red line) is defined by 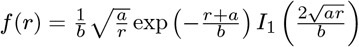, with *a* and 2*ab* being the mean and variance of the simulated data. The theoretical read number distribution is fit to the observed distribution with parameters *a* and *b*. This indicates that our simulation of pre growth is well approximated by the critical branching process with no fitness differences between lineages.

**Figure S5.**
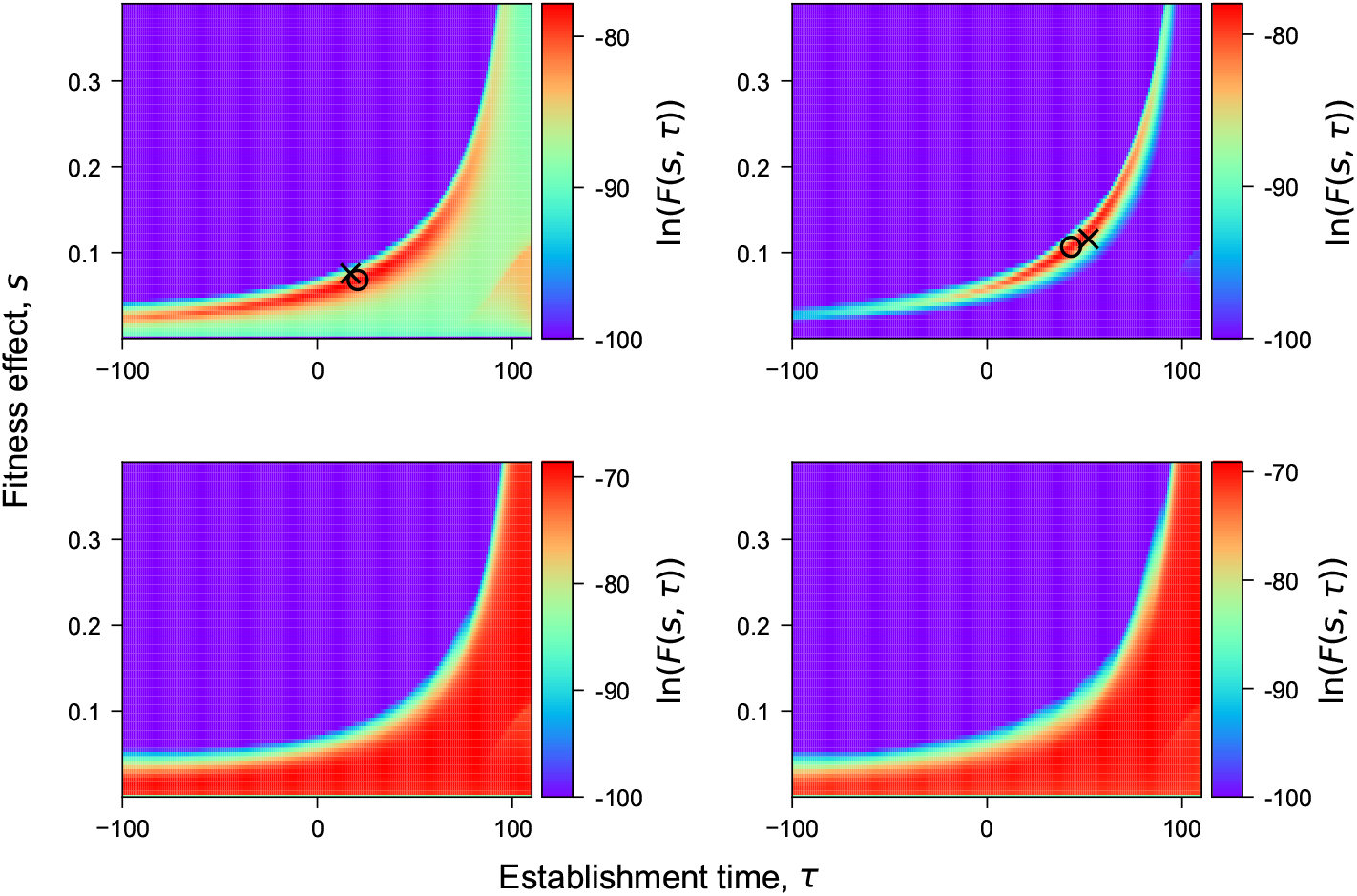
Optimization of the posterior likelihood. Value of the log-posterior likelihood ln*F*(*s,τ*) for two adaptive lineages on the 1st row, and two neutral lineages on the 2nd row, in the simulation from the 1st row and 4tcolumn of Figure 2. In the adaptive lineages, the fitness effect *s* and establishment time *τ* inferred by `FitMut2` is marked with ○. The true fitness effect *s*\* and occurrence time *t\** is marked with ×.

## Notes

https://github.com/FangfeiLi05/FitMut2

### Competing Interest Statement

The authors have declared no competing interest.

